# Genome assembly and annotation of the red flour beetle (*Tribolium castaneum)* from India

**DOI:** 10.1101/2024.05.20.594914

**Authors:** Shivansh Singhal, Chhavi Choudhary, Dipendra Nath Basu, Srijan Seal, Imroze Khan, Jayendra Nath Shukla, Deepa Agashe

## Abstract

The largest insect order, Coleoptera, includes several economically important beetles that also serve as major model species for biological research. Perhaps foremost among these is the red flour beetle, a global pest of stored grains and flour whose genome was sequenced in 2008. However, the currently available reference genome (Tcas5.2) is incomplete, fragmented and contains many gaps, and the Y chromosome is not assembled. Here we present inTcas1, an updated genome assembly and annotation of *T. castaneum* collected from India, assembled using both short and long read sequencing, and annotated using two transcriptome datasets. We report that inTcas1 has fewer gaps, less fragmentation, and many new genes and new isoforms of previously annotated genes. This new resource provides a useful update, comparison, and reference for new beetle genome assemblies. The first Y chromosome assembly for this species also provides critical data to study the evolution of insect sex chromosomes and sex determination systems.

**SIGNIFICANCE:** We present here an improved *T. castaneum* genome assembly (inTcas1) for beetles sampled from India, including the first Y chromosome of this important pest and laboratory model species. The new genome should facilitate more comprehensive analysis of Coleoptera genome and transcriptome datasets, especially beetle populations from Asia – the previously available genome, Tcas5.2, was assembled using the GA2 strain from USA. The updated genome should also facilitate analyses of genome evolution, including sex determination and sex chromosome dynamics; and the new gene annotations can expand the genetic toolkit for this beetle.

## INTRODUCTION

The red flour beetle *Tribolium castaneum* belongs to a basal insect lineage, the order Coleoptera (beetles), the most diverse taxon of the animal kingdom consisting of at least 350,000 species (Bouchard et al, 2017). Flour beetles are human associated pests that cause substantial economic losses via infestation of stored grains in warehouses and flour mills (Dyte & Blackman, 1970; Abdullahi et al, 2019). They have also been used for over a century as model species in ecology, evolutionary biology, and developmental biology (Sokoloff A. 1977; Pointer et al., 2021; Rösner et al. 2020). The *T. castaneum* genome was sequenced in 2008 (Richards et al, 2008), with subsequent corrections and additions (Herndon et al., 2020). More recently, several large datasets also report transcriptomes, RNAi screens, and chromatin profiles for *T. castaneum* (Rylee et al., 2018; Campbell et al., 2022), reinforcing its importance as a model system for biological research (Campbell et al., 2022).

However, genetic work with *T. castaneum* is still limited by the lack of a high-quality, complete reference genome. The published *T. castaneum* reference (Tcas5.2) was sequenced using Sanger and Illumina shotgun sequencing technology at 7X coverage (Richards et al, 2008). Despite subsequent updates (Herndon et al., 2020), 6.6% of the genome (13.5 Mb) consists of unspanned gaps and 8.3% (17 Mb) is in unplaced contigs (Volarić et al, 2022). The Y chromosome is also poorly assembled into 27 putative scaffolds. Large parts of the genome are likely missing in Tcas5.2, as evidenced by the inability to map sequencing reads from other studies (∼35% of reads from *T. castaneum* genomes from, India, Singhal et al, unpublished; and 16–87% of RNAseq reads from several transcriptome studies, https://www.ncbi.nlm.nih.gov/genome/annotation_euk/Tribolium_castaneum/103/). We present a new high quality contiguous *T. castaneum* genome (inTcas1) with ∼18X coverage, assembled using Illumina short read sequencing of DNA from 8 adults collected from India, and Oxford nanopore long reads from one female from an inbred population (Table S1, S2). We assembled draft genomes for all individuals, filled gaps, and rescaffolded the genome, allowing us to fill 90% of the gaps in Tcas5.2 and add 13 Mbp of new sequence.

## RESULTS AND DISCUSSION

### Improved genome assembly and annotation

The previously published *T. castaneum* genome (Tcas5.2) contains many gaps, and our main objective was to address this problem (Richards et al, 2008). We generated contigs by stitching together our Illumina short reads, assembled contigs from one beetle into scaffolds using Tcas5.2 linkage groups (LGs) as reference, and filled gaps in scaffolds using 6 other draft genomes as well as corrected Nanopore long reads (Figure S1). We estimated the genome size with GenomeScope2.0 as 179–189 Mbp; our final reference assembly (inTcas1) had a slightly smaller genome size of 169.11 Mbp. The new genome represents a substantial improvement over Tcas5.2, with fewer gaps and scaffolds, less fragmentation (e.g., higher N50 values, Table 1), substantial amount of new sequence data (Table S3), and 27 Mb of unplaced contigs (Table S4). We added new protein coding genes and isomers of existing genes on all linkage groups, with 22,171 new gene models relative to Tcas5.2. Excluding new isoforms of previously annotated genes, we report 979 new non-overlapping non-redundant gene models in inTcas1 (Table 1, Table S4). The inTcas1 genome also has more pseudogenes, tRNA genes, and non-coding RNAs (Table 1). The BUSCO insect database for benchmarked universal single copy orthologs (BUSCOs) indicates that the inTcas1 genome is 98.3% complete (Table S5).

**TABLE 1:** Genome assembly and annotation summary. Genome quality comparison for *Tribolium castaneum* genome Tcas5.2 (Richards et al, 2008) and inTcas1 (this study).

|  | **Tcas5.2** | **inTcas1** |
| --- | --- | --- |
| Size (Mbp) | 165.94 | 169.11 |
| Ungapped Length (Mbp) | 152.43 | 165.9 |
| Scaffold N50 (Mbp) | 4.46 | 14.47 |
| Number of Ns (Mbp) | 13.52 | 3.21 |
| Number of Scaffolds | 2149 | 1576 |
| Scaffold L50 | 12 | 5 |
| Number of ncRNA | 460 | 764 |
| Genes (non-overlapping) | 5670 | 6649 |
| Genes (isoforms) | 14491 | 36233 |
| Pseudogenes | 9 | 34 |
| rRNA genes | 2 | 2 |
| tRNA genes | 79 | 150 |

### Structural differences between Tcas5.2, inTcas1, and sister species

The broad structure of Tcas5.2 and inTcas1 is consistent, as observed in the proportion of locally co-linear blocks (LCBs) shared between both assemblies, with two exceptions. Some LCBs from Tcas5.2 LGX could not be assembled into linkage groups in inTcas1, while others from Tcas5.2 LG3 are now assembled into inTcas1 LGX (Figure 1A; Figure S2). Translocations between autosomes and X chromosome are not surprising, and were previously observed in other beetle species (Bracewell et al, 2023). Broadly, the discrepancies between Tcas5.2 and inTcas1 could reflect divergence between *T. castaneum* populations from USA and India respectively, or arise due to incorrect scaffolding in Tcas5.2. The differences are unlikely to reflect mis-assembly in inTcas1 (because we confirmed true structural differences using misassembly correction, SI methods), enrichment for repeat regions (only ∼34% of the LCBs consists of repeats), or enrichment for new sequences that were not present in Tcas5.2 (only ∼13% of the LCBs are new). The inTcas1 assembly also has several smaller sequence additions, deletions, inversions, and translocations (within and across linkage groups), but these are distributed all over the genome (Figure 1B; Figure S2).

**Figure 1:**
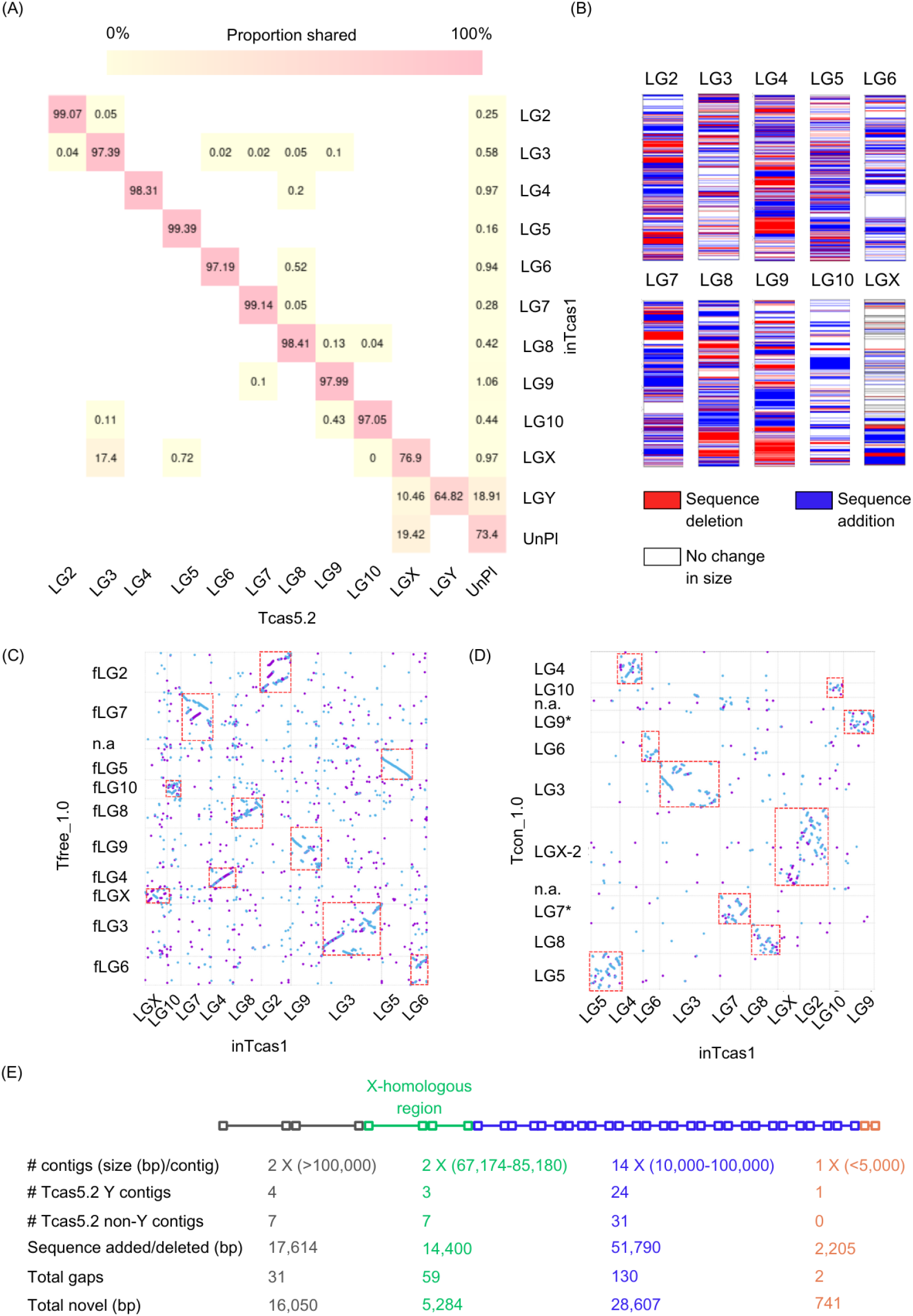
Comparing inTcas1 with Tcas5.2 and closely related species. (A) Heatmap showing the proportion of the total length of ungapped chromosomes of inTcas1 LGs (on the y-axis) that matches with Tcas5.2 LGs (on the x-axis). Matching refers to >95% sequence identity and >60% coverage of target locally colinear blocks (LCBs). For example, 99.07% of inTcas1 LG2 matches with Tcas5.2 LG2. UnPl=unplaced contigs. (B) Schematic illustrating the location of sequence additions (blue) and deletions (red) in inTcas1 relative to Tcas5.2. The size of colored regions is not proportional to the addition/deletion size, but instead indicates the size of the LCB in which the addition/deletion was found. (C–D) Comparison of inTcas1 (x-axis) with (C) *T. freemani* and (D) *T. confusum*. Each dot represents an LCB (defined as above), colored to indicate its relative orientation in the two genomes (purple=same, blue=opposite). n.a. indicates unplaced contigs, and asterisks indicate LGs that were not named as chromosomes in the reference genome database. Red boxes highlight matching LGs in the two genomes. (E) Schematic of the inTcas1 Y chromosome indicating 19 contigs and homology with LGX, and a summary of comparison with Tcas5.2 sequences on putative Y scaffolds and elsewhere. Contigs are colored by size bins, as indicated.

Comparing inTcas1 with the closest relatives *T. freemani* (Volarić et al, 2022) and *T. confusum* (Bracewell et al, 2023), we observed that the former shows better synteny between linkage groups (Figures 1C-D), consistent with the recent finding that *T. freemani* is the sister species of *T. castaneum*, instead of *T. confusum* as was previously believed (Ramesh et al 2021). However, across the ∼13.7 My (million years) divergence between *T. castaneum* and *T. freemani*, we do see several structural differences within LGs. The *T. confusum* X chromosome shows synteny with both LGX and LG2 of inTcas1, supporting the recent finding that the *T. confusum* X chromosome has fused with the ancestral LG2 (Bracewell et al, 2023). Thus, despite improved assembly, these broad patterns of genome evolution across the genus *Tribolium* do not change.

### Scaffold level assembly of Y chromosome

Using differences in kmers and their frequencies between males and females (Figure S3), we assembled the first reported *T. castaneum* Y chromosome of size 0.8 Mbp, with 6.27% new sequences (Figure 1E, Table S3, Table S4). Of the 38 genes (Table S4), half were also present in scaffolds labelled as Y in the Tcas5.2. Overall, a large proportion of the Y chromosome (∼65%) represents unplaced contigs that were labelled as Y in Tcas5.2 (Y contigs), ∼19% was in unplaced contigs that were not labelled as Y (non-Y contigs), and ∼10% was homologous to LGX (Figures 1A and 1E). Together, these results indicate that Y chromosome assembly rather than sequencing was a limiting factor in Tcas5.2. We hope that the improved *T. castaneum* genome will facilitate future genetic work with this important species.

## METHODS

### Sample collection, genomic DNA extraction and sequencing

In 2013, we set up laboratory stocks of three *Tribolium castaneum* populations collected from grain storehouses across India (populations 12, 13, 18, with ∼50–100 founders each), and one inbred population from a packet of infested commercially available flour in Bengaluru (population 1, with a single founding female) (Table S1 and Table S2). Stocks were maintained at 33±1 °C in commercially available whole wheat flour without any supplements, with large population sizes (∼2000 adults) maintained on a 4 to 5–week discrete generation cycle. Fresh flour was supplied at each generation, or if stocks were overcrowded, ∼3 weeks after initiation.

We used individual beetles from these stocks for genome sequencing. For short read sequencing, we extracted DNA from 8 individuals using the Promega Wizard genomic DNA extraction kit with the default protocol for tissue DNA (except an overnight proteinase K treatment and 3× wash with 70% ethanol), and sequenced it using Illumina paired end short read sequencing (2×150bp). For long read sequencing, we imposed inbreeding in a new lineage derived from population 1, allowing a single pair of siblings to mate in each generation for 4 generations. We used one female from the fifth generation to extract high molecular weight DNA (Volarić et al, 2022) using the Qiagen DNeasy blood and tissue kit, confirmed that the peak fragment size was >1Kb, and sequenced it on the Oxford Nanopore MinION platform.

### *De novo* genome assembly and reference-guided genome scaffolding

We followed several iterative steps to generate our reference genome, described briefly here (Figure S1; detailed in supplementary methods). From the >15 million read pairs for each individual from short read sequencing (Table S1), we removed low quality bases and estimated the frequency of all kmers using jellyfish (Marcais and Kingsford, 2011). We mapped the reads to Tcas5.2, but found that ∼32% of our reads did not align to the reference (range 24-39% across samples). We assembled short reads into contigs using SPAdes (Prjibelski et al, 2020), polished it and corrected mis-assemblies using Pilon (Walker et al, 2014), and removed haplotigs using Purge Haplotigs v1.0.0 (Roach et al, 2018). We used one genome (1A) to scaffold into chromosomes using Tcas5.2 (Accession: GCF_000002335.3) as the reference using RagTag (Alonge, M, 2021; Alonge et al, 2022). Then we used 6 other draft genomes and long reads to fill or extend gaps in genome 1A using TGS-GapCloser (Xu et al, 2020) (Table S6).

To assemble the Y chromosome with short read data for all samples, we used DiscoverY (Rangavittal et al, 2019) to identify male-specific kmers. We scaffolded them using contigs labelled as Y in Tcas5.2 (Figure S3), removing redundant contigs present in the assembled autosomes to obtain a putative Y chromosome. We added this Y chromosome sequence to the genome assembled in previous steps, to obtain the complete nuclear genome of the red flour beetle (inTcas1). The unplaced contigs were combined into a single scaffold by adding 50 placeholder Ns between contigs. We confirmed that there was no bacterial (or other) contamination in inTcas1 using the FCS-GX NCBI tool with the entire database (Astashyn et al, 2023). We calculated basic assembly metrics using QUAST (Gurevich et al, 2013) and GAEP (Zhang et al, 2023). We estimated the completeness of the genome using BUSCO with odb10-insecta (Simao et al, 2015; database obtained on 10 December 2021). For chromosome level comparisons between inTcas1 and Tcas5.2 (Accession: GCF_000002335.3), Tfree1.0 (Accession: GCA_939628115.1) and Tcon1.0 (Accession: GCA_019155225.1), we used nucmer, MUMmer and SibiliaZ (Kurtz et al, 2004; Delcher et al, 2002; Minkin and Medvedev, 2020).

### Genome annotation

To find and annotate genomic features in inTcas1, we used maker2.0 (Cantarel et al, 2008), which incorporates EST evidence (e.g., from transcriptomes), a custom repeat library, *ab initio* gene prediction and homology-based gene prediction. We assembled a transcriptome using two RNAseq datasets sequenced on the Illumina platform (150x2 PE): dataset 1 comprising 1.7 million reads from 12-day old females from an outbred laboratory population of *T. castaneum*, and dataset 2 comprising 4.1 million reads from males, females, and eggs from a single wild-collected population. We assembled the two datasets using SOAPdenovo-Trans (Boetzer and Pirovano, 2012) (Table S7). We constructed a *de novo* species specific repeat library using RepeatModeller v1.0.7 (Flynn et al, 2020) (Table S8). We used AUGUSTUS (Stanke and Morgenstern, 2005) and Exonerate (Slater and Birney, 2005) for protein homology-based gene search, SNAP (Korf I, 2004) and GlimmerHMM (Majoros et al,2004) for *ab initio* gene search, and Liftoff (Shumate and Salzberg, 2021) to transfer annotations from Tcas5.2. All the evidence above was then used to identify putative genomic features such as genes, exons, CDS, and some types of RNA and repeats in the Maker annotation pipeline (Holt and Yandell, 2011), run three times iteratively. After the third round we found >95% of all genes had AED scores (annotation edit distance) of <0.5 which is ideal for calling genes (Figure S4, Table S9). We removed overlapping and redundant gene models based on coordinates and AED scores using custom python and bash scripts. We then combined the annotation from Maker and Liftoff to create a final genome features file containing annotations, without duplicate annotations from different tools.

## DATA AVAILABILITY

Raw reads for all 8 draft genomes and 2 transcriptome datasets, genome and annotations are available under NCBI BioProject ID PRJNA1077124. Scripts related to this project are available on GitHub (https://github.com/shivanshss/tcas_india_genome_assembly).

## ACKNOWLEDGEMENTS

We thank Pratibha Sanjenbam for critical comments on the manuscript; Aparna Agarwal for DNA extraction for short read sequencing; Sneha Garge, Awadhesh Pandit and the NCBS sequencing facility for help with long read sequencing. We thank the NCBS High Performance Computing (HPC) facility and Aswin Seshasayee for access to computing clusters. We acknowledge funding and support from the National Centre for Biological Sciences (NCBS–TIFR) and the Department of Atomic Energy, Government of India (Project Identification No. RTI 4006) to DA, a SERB Women Research Excellence award to DA (WEA/2020/000030), a DBT/Wellcome Trust India Alliance Intermediate Fellowship (IA/I/20/1/504930) to IK, a SERB-DST grant to (ECR/2017/003370) to IK, and a DST-SERB grant (EMR/2017/001378/AS) to JNS.

## AUTHOR CONTRIBUTIONS

S Singhal and DA conceived the project and designed the work; S Singhal, CC, S Seal and DNB collected sequencing data; S Singhal analysed data; DA, JNS and IK directed the project; S Singhal and DA wrote the manuscript; DA, IK and JNS acquired funding.

