## Supplementary material for "Genome assembly and annotation of the red flour beetle (*Tribolium castaneum)* from India"

### SUPPLEMENTARY METHODS

#### De novo genome assembly and reference-guided genome scaffolding

We sequenced DNA from a total of 9 individuals – 8 individuals for short read sequencing (Table S1) and 1 inbred female for long read sequencing (Table S2). For short read data, we used FastQC v.0.11.8 (Andrews et al, 2010) to visualize and obtain a summary of read quality of the raw sequence data and trimmed or masked low-quality bases using Trimmomatic v.0.38 (Bolger et al, 2014). We obtained 16 to 30 million reads for 8 samples (Table S1). We used jellyfish to compute kmer frequencies (Marcais and Kingsford, 2011). We then used GenomeScope2.0 with default parameters (Ranallo-Benavidez et al, 2020) to estimate the genome size and quantify heterozygosity using the frequencies of unique kmers (non-overlapping set of kmers). For validation, we used kmc to ascertain correct estimation of genome size and kmer frequencies (Kokot et al, 2017).

We first tried to map our short reads to the published *T. castaneum* reference genome (Tcas5.2), using the bwa mem algorithm (Li H. 2013). However, 24-39% of reads were not aligned. Hence, we assembled Illumina short reads *de novo* separately for each of the 8 samples using SPAdes ver3.15.2 (Prjibelski et al, 2020) to form contigs. We used Purge Haplotigs v1.0.0 (Roach et al, 2018) to remove redundant contigs from each assembly, identified using the coverage histogram (showing the frequency of unique kmers) given by Jellyfish for each population (e.g., for individual 1A, we chose parameter values -l 5 -m 30 -h 190 based on its kmer coverage histogram from GenomeScope). Purge Haplotigs uses read depth coverage to reduce over-purging of repetitive regions and paralogous contigs in highly heterozygous assemblies. We set the threshold for identifying a contig as a haplotig (a contig with very high similarity to another contig) to -a 40, which is lower than the default setting, since we observed high heterozygosity in our short read sequences. At the end of these steps, we obtained draft genome assemblies for each of our 8 samples. Next, we used short reads for each sample and respective draft genomes to correct mis-assemblies using three rounds of Pilon genome polishing (Walker et al, 2014). A final round of Purge Haplotigs was then performed to reduce duplication further, using more stringent parameter values (e.g., for individual 1A, -l 10 -m 50 -h 150', and for all samples, the haplotig threshold as -a 80).

Ultimately, our goal was to produce a single high quality reference genome. We therefore started scaffolding with one of the most inbred female genome (1A), and used its draft assembly to scaffold into pseudo-chromosomes matching Tcas5.2 with the 'scaffold' feature of RagTag (Alonge, Michael, et al, 2021; Alonge et al, 2022). For validation, we used MeDuSa (Bosi et al, 2015) and RAILS (Warren R. L., 2016) for reference-based scaffolding. There was no difference in sizes and identity of contigs placed on linkage groups between scaffolding tools. We then filled gaps in this genome using the draft assemblies from the 6 other individuals as well as long reads from the hyper-inbred individual of population 1 (corrected using canu (Table S2)) using TGS-gapCloser (Xu et al, 2020) (Table S6). Two draft genomes (1B and 18B) were not used for gapfilling due to low size and high fragmentation. For validation, we used GapCloser (Luo et al, 2012), LR-Gapcloser (Xu et al, 2019) and GapFiller (Boetzer and Pirovano, 2012) to determine the tool with best gapfilling metrics. We then merged all short reads from all individuals, and performed one more round of Pilon polishing using short reads from all samples.

For the final assembled genome (inTcas1), we calculated basic assembly statistics using QUAST (Gurevich et al, 2013) and GAEP (Zhang et al, 2023). We estimated the completeness of the genome using BUSCO with odb10-insecta (Simao et al, 2015; database obtained on 10 December 2021). For chromosome level comparisons between Tcas5.2

(Richards et al, 2008), inTcas1 (this study), Tcon1.1 (Bracewell et al, 2023) and Tfree1.1 (Volarić et al, 2022) we used nucmer, MUMmer and SibiliaZ (Kurtz et al, 2004; Delcher et al, 2002; Minkin and Medvedev, 2020).

**Y chromosome assembly**

For short read data for 6 males and 2 females, we used DiscoverY (Rangavittal et al, 2019, BMC genomics) to identify male-specific kmers that had half the average male autosomal coverage. For validation, we used findZX (Sigeman et al, 2022), finding the same male specific contigs.

Several kmers can be assigned to a single contig, and therefore, the proportion of male-specific kmers in each contig (here, the threshold was set to 0.2) was used to identify male-specific contigs. In this way, we identified contigs that likely belong to the Y chromosome. We then filtered the contigs based on presence in atleast 2 other male samples. We used these filtered contigs to assemble a Y chromosome using SPAdes ver3.15.2 (Prjibelski et al, 2020) and scaffold it using RagTag scaffold using the contigs labelled as putative Y-chromosome in Tcas5.2. To make sure that Y scaffolds are not represented twice in the assembly, we aligned scaffolds from Y to all autosomes of our new genome inTcas1 (excluding the X chromosome because there is known homology between X and Y chromosomes in some species, Cortez et al, 2014; Blackmon et al, 2017). To filter contigs that were erroneously assigned to the Y chromosome, we removed contigs with >80% identity with an autosomal contig, and an aligned length to query length ratio >0.6. We used BLASTn in custom bash scripts for this step. After 2 iterations of Pilon error polishing, we obtained a Y chromosome in 19 pieces with some unspanned gaps. We added this Y chromosome sequence to the genome assembled in previous steps, to obtain the complete nuclear genome of the red flour beetle. This genome contains 9 autosomes, one X chromosome, one Y chromosome and concatenated unplaced contigs. The unplaced contigs were combined into a single scaffold by adding 50 placeholder Ns between contigs for annotation step.

**Genome annotation**

To find and annotate genomic features in the new genome, we used maker2.0 (Cantarel et al, 2008), which incorporates evidence from transcriptomes, a custom repeat library, ab initio gene prediction. We assembled a transcriptome for EST evidence using two transcriptome datasets. For the first dataset, we crushed ten 2-day-old adult females beetles from an outbred population together, and extracted RNA from using Qiagen RNeasy kit. This outbred population was created using individuals from 12 populations of *T. castaneum* from India. We sequenced this RNA using Illumina Novaseq 6000 PE150 technology and obtained 17.5 million 150bp paired-end reads after trimming. For the second RNA dataset, we extracted RNA from 6 days old adults (5 males and 5 females) and pooled 0-6 h old eggs (~80 to 100) using Trizol RNA extraction protocol. These individuals were obtained from Entomology department of Punjab Agricultural University, originally collected from local natural populations of *Tribolium castaneum* from India. We pooled RNA in equimolar proportion from 5 adults, males and females taken separately. We sequenced these three samples of pooled RNA (males, females and eggs) using Illumina HiSeq PE150 technology and obtained >4 million 150bp paired end reads.

We removed reads of size <50 bp and trimmed or marked (as N) bases that had a quality score <20 using fastQC and Trimmomatic (Andrews et al, 2010; Bolger et al, 2014). We used these reads to assemble the transcriptome using SOAP-denovoTrans (Boetzer and Pirovano, 2012) for kmer values ranging from 21 to 31 in steps of 2 and 41 to 121 in steps of

10. We chose the best assembly based on total assembly length and fragmentation level (K=21 for dataset 1 and K=31 for dataset 2) (Table S7). We used assembled transcriptome as EST evidence for the annotation pipeline.

Next, we constructed a de novo species specific repeat library using RepeatModeller v1.0.7 (Flynn et al, 2020) to identify repeats, and removed potential protein-coding genes by searching against the GenBank non-redundant (nr) protein database for Arthropoda (e value <10-3) using Blastx. We added known transposons from RepBase (Jurka et al, 2005) to create the final custom repeats library. To mask repeats and estimate the abundances of all predicted repeats in the genome assembly, we used RepeatMasker v4.0.7 (Nishimura, D., 2000) (Table S8). Finally, we used AUGUSTUS (Stanke and Morgenstern, 2005) and Exonerate (Slater and Birney, 2005) for protein homology-based annotation, SNAP (Korf I, 2004) and GlimmerHMM (Majoros et al,2004) for ab initio gene finding based on known structural features of genes (e.g., ORFs and transcription factor binding sites), and Liftoff (Shumate and Salzberg, 2021) to lift over annotations from Tcas5.2.

All the evidence above was then used to annotate one chromosome at a time, with all unplaced contigs concatenated and treated as a single chromosome. We removed overlapping and redundant gene models based on genome coordinates and annotation confidence scores using a custom python script. We used default parameters of the maker annotation pipeline (Holt and Yandell, 2011) to identify putative genomic features such as genes, exons, CDS, and some types of RNA and repeats (Table S9). We then combined the annotation from maker and liftoff to create a final genome features file containing annotations without duplicate annotations from different tools.

Genes were identified as pseudogenes if two very similar copies of the same gene were found with one copy showing at least one of the following features: missing promoter, disrupted start or stop codon, large deletion, missing intron or frameshift. We manually checked for features such as the location of the pair of putative pseudogenes, estimated break points, gene size, and % identity with the closest paralog, and removed gene pairs that were overlapping or were present on separate linkage groups (these are most likely homologous regions that were incorrectly identified as pseudogenes).

### SUPPLEMENTARY FIGURES

**Figure S1: Genome assembly steps**. Schematic showing the process of genome assembly from short and long reads along with tools used at each step. Tools marked in black were used in this analysis and those in purple were used for validation at various steps. The focus of this pipeline was to fill gaps and produce a complete, ungapped sequence of linkage groups.

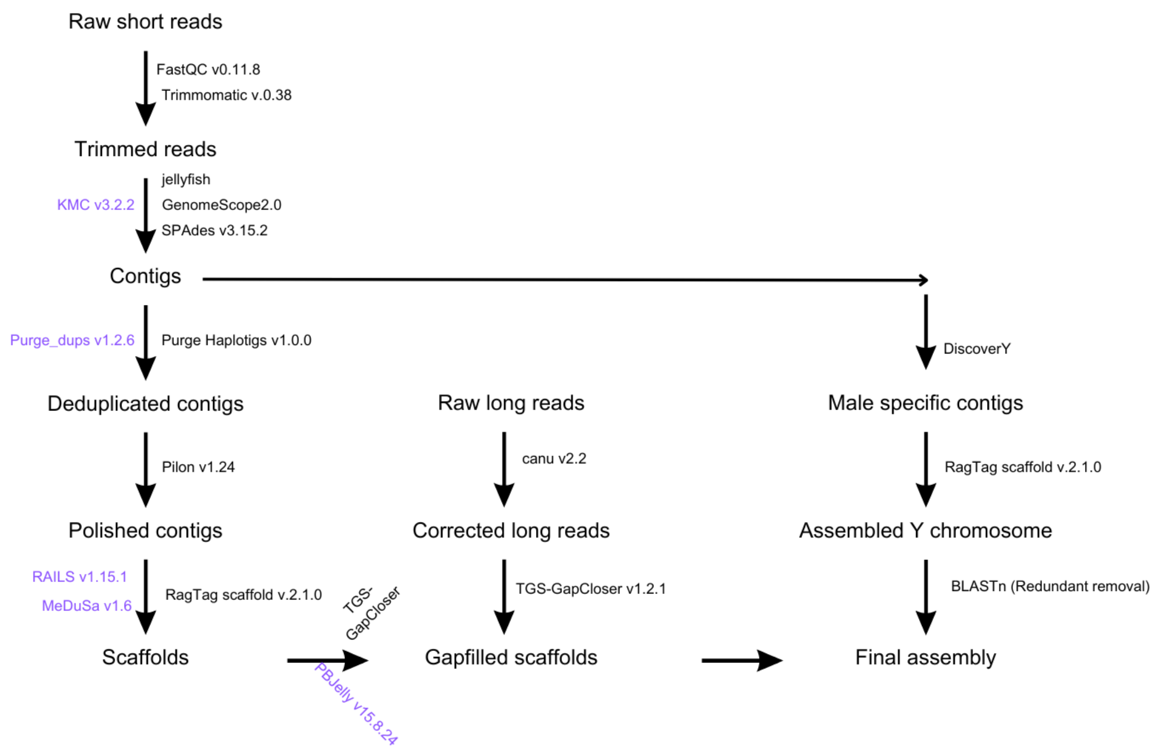

**Figure S2: Comparison of inTcas1 and Tcas5.2.** Dotplot comparing inTcas1 (x-axis) with Tcas5.2 (y-axis). Each dot represents an LCB (defined as blocks of sequence in inTcas1with >95% sequence identity and >60% coverage of sequence in Tcas5.2); colors indicate its relative orientation in the two genomes (purple=same, blue=opposite). Some unplaced contigs (UnPl) could be placed on various linkage groups and are indicated in red. Black boxes indicate best-matched LGs across the two genomes.

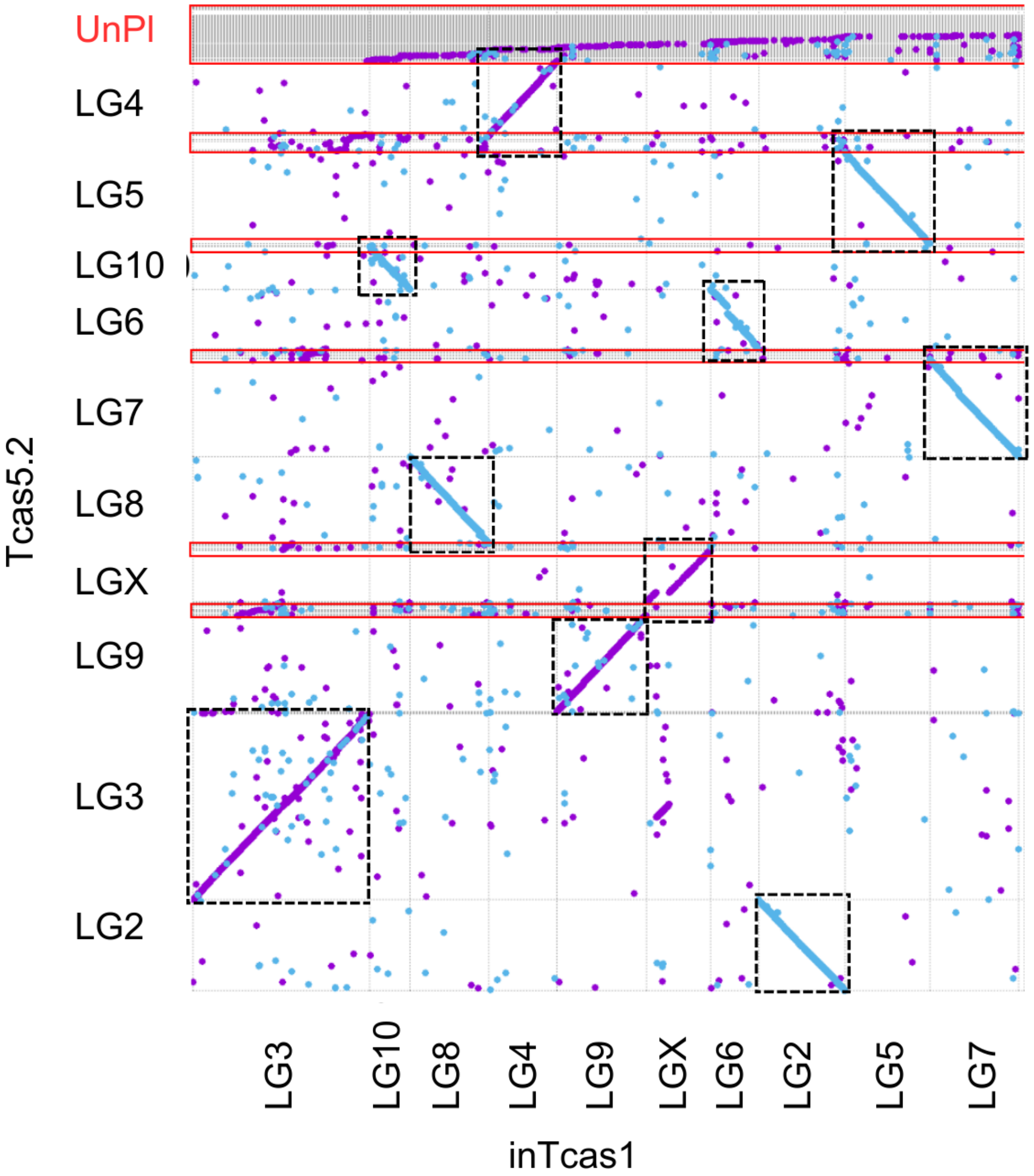

**Figure S3: Identifying male-specific contigs.** Scatterplot showing the proportion of a male contig (indicated by a point) shared with contigs from females (x-axis), and the corresponding coverage of that contig in males (y-axis). Panel labels indicate the identity of male individuals. Male contigs are likely to be those that share a lower fraction of the female contigs, and have half the autosomal coverage. Such putative male contigs (marked in green) are then scaffolded into the Y chromosome.

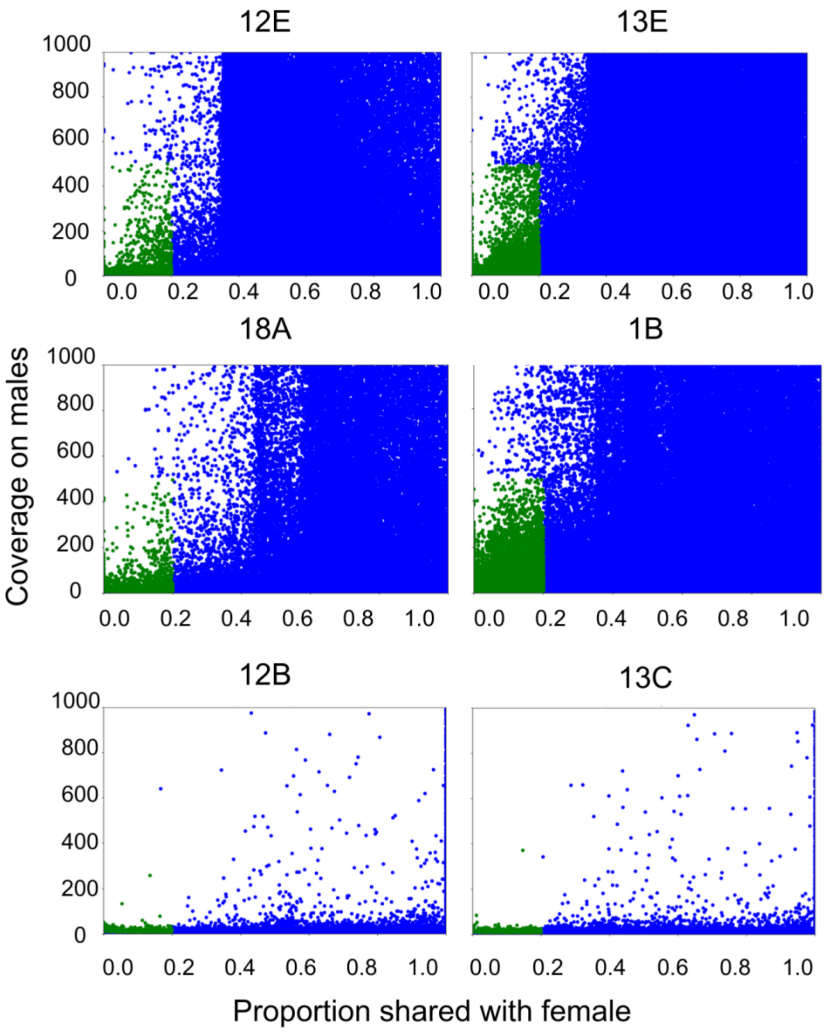

**Figure S4: Confidence in gene annotation.** Annotation edit distance (AED) is a maker score given to every gene that is predicted by the maker tool; lower AED scores indicate higher confidence in the annotation. >95% genes have AED scores <0.5 (threshold for confident gene annotation).

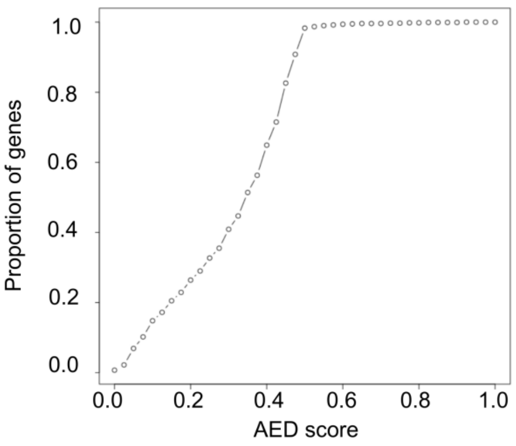

**SUPPLEMENTARY TABLES**

**Table S1: Details of beetle samples and summary of short read sequencing.** Short reads were obtained for 2-3 individuals from 4 different populations (locations) across India. QC includes quality estimation and trimming or masking of low-quality sequences.

| Population and ID | Sex | Location | Number of reads before QC (million) | Number of reads after QC (million) | Read size before QC | Read size after QC |
|---|---|---|---|---|---|---|
| 1A | F | Bengaluru | 25.48 | 23.85 | 150 | 142-150 |
| 1B | M | Bengaluru | 23.73 | 22.42 | 150 | 121-138 |
| 12D | F | Puducherry | 33.06 | 30.87 | 150 | 141-150 |
| 12B | M | Puducherry | 23.47 | 21.89 | 150 | 114-129 |
| 12E | M | Puducherry | 23.60 | 21.99 | 150 | 122-127 |
| 13C | M | Indore | 17.52 | 16.3 | 150 | 120-130 |
| 13E | M | Indore | 24.99 | 23.34 | 150 | 134-136 |
| 18A | M | Chandigarh | 25.68 | 24.54 | 150 | 114-120 |
| 18B | M | Chandigarh | 20.55 | 19.12 | 150 | 124-130 |

**Table S2: Long read summary.** Long reads were obtained by sequencing a single adult female from a hyper inbred line of beetles, using Oxford Nanopore long read sequencing technology. ONT 1 and ONT 2 are 2 sets of raw long reads. ONT corrected are merged and quality corrected reads from ONT 1 and ONT2.

| ID | Total size in basepairs (million) | Number of reads (thousand) | Minimum read size | Maximum read size |
|---|---|---|---|---|
| ONT 1 | 539.384 | 309.043 | 3 | 89,747 |
| ONT 2 | 763.110 | 473.810 | 3 | 82,401 |
| ONT corrected | 506.849 | 336.989 | 200 | 35,408 |

**Table S3: Percent of putatively new regions on every chromosome in inTcas1 compared to Tcas5.2.** Proportion (by length) of locally collinear blocks (LCBs) in linkage groups of inTcas1 that are new relative to Tcas5.2 (Regions of inTcas1 not find in Tcas5.2). Proportions are calculated by dividing the sum of length of all regions not found Tcas5.2 to the total ungapped length of the inTcas1 LG.

| | **Percent novel in inTcas1** |
|---|---|
| LG2 | 0.62 |
| LG3 | 1.7 |
| LG4 | 0.51 |
| LG5 | 0.45 |
| LG6 | 1.3 |
| LG7 | 0.53 |
| LG8 | 0.99 |
| LG9 | 0.85 |
| LG10 | 1.97 |
| LGX | 4.01 |
| LGY | 6.27 |
| unplaced | 7.17 |

**Table S4: Chromosome-wise assembly and annotation summary.** Unpl stands for unplaced contigs.

| | LG2 | LG3 | LG4 | LG5 | LG6 | LG7 | LG8 | LG9 | LG10 | LGX | LGY | Unpl |
|---|---|---|---|---|---|---|---|---|---|---|---|---|
| Size (Tcas5.2) (mbp) | 15.26 | 31.38 | 12.29 | 15.45 | 10.08 | 16.48 | 14.58 | 16.18 | 7.22 | 8.67 | 0.88 | 17.42 |
| Size (inTcas1) (mbp) | 14.80 | 30.16 | 11.56 | 14.47 | 8.20 | 15.27 | 13.56 | 15.24 | 6.90 | 11.00 | 0.856 | 27.05 |
| Ungapped length (Tcas5.2) (mbp) | 14.56 | 29.60 | 11.63 | 14.34 | 7.98 | 15.19 | 13.62 | 15.01 | 6.78 | 7.42 | 0.69 | 15.59 |
| Ungapped length (inTcas1) (mbp) | 14.6 | 29.68 | 11.45 | 14.30 | 8.04 | 15.07 | 13.34 | 15.02 | 6.77 | 9.67 | 0.833 | 27.05 |
| Number of Ns (Tcas5.2) (mbp) | 0.709 | 1.77 | 0.658 | 1.12 | 2.10 | 1.29 | 0.963 | 1.17 | 0.438 | 1.24 | 0.188634 | 1.83 |
| Number of Ns (inTcas1) (mbp) | 0.179 | 0.471 | 0.112 | 0.165 | 0.161 | 0.194 | 0.219 | 0.221 | 0.135 | 1.267 | 0.023 | 0 |
| Ns filled | 0.53 | 1.299 | 0.546 | 0.955 | 1.939 | 1.096 | 0.744 | 0.949 | 0.303 | -0.027 | 0.157 | NA |
| nc_rna | 54 | 44 | 82 | 54 | 0 | 96 | 42 | 32 | 28 | 28 | NA | NA |

| | | | | | | | | | | | | |
|---|---|---|---|---|---|---|---|---|---|---|---|---|
| (Tcas5.2) | | | | | | | | | | | | |
| nc_rna (inTcas1) | 72 | 152 | 24 | 47 | 52 | 56 | 55 | 70 | 7 | 55 | 26 | 148 |
| Genes (non-overlapping) (Tcas5.2) | 546 | 360 | 1063 | 676 | 1 | 1044 | 421 | 421 | 379 | 439 | NA | NA |
| Genes (non-overlapping) (inTcas1) | 719 | 728 | 302 | 879 | 330 | 951 | 968 | 365 | 222 | 364 | 28 | 793 |
| Genes (all) (inTcas1) | 2762 | 3561 | 2523 | 3050 | 1661 | 3283 | 2922 | 2442 | 1027 | 1304 | 38 | 2139 |

**Table S5: BUSCO summary.** We used the *Insecta* database with 1367 expected BUSCOs. New genome inTcas1 is 98.32% complete with 0.4% duplication.

| **BUSCO** | **Number of genes** | **Percentage of total** |
|---|---|---|
| Total BUSCO groups searched (T) | 1367 | |
| Complete BUSCO (C) | 1344 | 98.32 |
| Complete and single-copy BUSCO (S) | 1338 | 97.88 |
| Complete and duplicated BUSCO (D) | 6 | 0.44 |
| Fragmented BUSCO (F) | 18 | 1.32 |
| Missing BUSCO (M) | 5 | 0.37 |

**Table S6: Gapfill metrics:** Genome 1A was used as the base genome onto which gap filling was done using 6 draft genomes and one long read dataset. Filling steps 5, 6 and 7 show a decrease in ungapped length as more and more gaps could be fully spanned. Number of Ns decrease through all the steps.

| | **Fill 1** | **Fill 2** | **Fill 3** | **Fill 4** | **Fill 5** | **Fill 6** | **Fill 7** |
|---|---|---|---|---|---|---|---|
| Query draft genome | 12B | 12D | 12E | 13C | 13E | 18A | ONT corrected |
| Ns filled in kbp | 660.926 | 1233.746 | 98.499 | 3.288 | 37.163 | 79.835 | 415.844 |
| Size in Mbp | 130.200 | 130.296 | 130.068 | 130.046 | 129.922 | 129.906 | 130.200 |
| Ungapped Length (Mbp) | 126.904 | 126.961 | 127.973 | 128.051 | 127.951 | 127.931 | 127.943 |
| Number of Ns (Mbp) | 2.635 | 2.101 | 1.995 | 1.991 | 1.934 | 1.895 | 1.841 |

**Table S7: Summary of transcriptome assembly.** K denotes the length of kmers used for assembly. We used kmer lengths with highest total size and number of scaffolds (highlighted in bold) as EST evidence for maker pipeline.

| Kmer_size | Total size | Number of scaffolds | Mean size | Median size | Longest sequence length |
|---|---|---|---|---|---|
| **Dataset 1** | | | | | |
| **K21** | **20,978,101** | **46,128** | **454** | **237** | **12636** |
| K23 | 20,637,306 | 46,667 | 442 | 231 | 13397 |
| K25 | 20,264,272 | 48,143 | 420 | 226 | 11042 |
| K27 | 19,860,640 | 49,927 | 397 | 217 | 7249 |
| K29 | 19,375,188 | 52,155 | 371 | 208 | 12077 |
| K31 | 18,730,403 | 54,685 | 342 | 198 | 8609 |
| **Dataset 2** | | | | | |
| K21 | 45,862,859 | 133,919 | 342 | 205 | 12833 |
| K23 | 47,062,578 | 139,203 | 338 | 203 | 12504 |
| K25 | 48,063,810 | 146,000 | 329 | 198 | 14905 |
| K27 | 49,234,156 | | 316 | 189 | 12833 |
| K29 | 50,458,214 | 165,552 | 304 | 179 | 9666 |
| **K31** | **51,494,852** | **174,006** | **295** | **172** | **12833** |

**Table S8: Comparison of numbers of repeat elements in Tcas5.2 and inTcas1.** DNA repeat elements (DNA), long interspersed nuclear elements (LINE), short interspersed nuclear element (SINE) Rolling circle repeat (RC), RNA repeats (rRNA, tRNA) and long terminal repeats (LTR). Repeat types and numbers in Tcas5.2 are taken from Volarić et al, 2022.

| | **Tcas5.2** | **inTcas1** |
|---|---|---|
| DNA (X1000) | 6.51 | 9.45 |
| LINE (X1000) | 3.18 | 2.41 |
| LTR (X1000) | 0.83 | 1.13 |
| RC (X1000) | 0.67 | 0.97 |
| SINE (X1000) | 0.24 | 0.69 |
| Total % of genome by length | 64 | 69 |

**Table S9: Annotation summary.** Number of gene models (including overlaps) with AED score below 0.5 found after 3 rounds of maker by each tool used for gene finding.

| Tool | Number of annotated elements |
|---|---|
| AUGUSTUS | 14,427 |
| BLAST (BLASTn, BLASTx, tBLASTx) | 171,340 |
| cdna2genome | 20,785 |
| maker | 160,035 |
| liftoff | 14,096 |
| protein2genome | 65,974 |
| est2genome | 218033 |
| Repeats (Repeatrunner, repeatMasker) | 110,364 |
